# Repetition Attenuates the Influence of Recency on Recognition Memory: Behavioral and Electrophysiological Evidence

**DOI:** 10.1101/826693

**Authors:** John E. Scofield, Mason H. Price, Angélica Flores, Edgar C. Merkle, Jeffrey D. Johnson

## Abstract

Studies of recognition memory often demonstrate a recency effect on behavioral performance, whereby response times (RTs) are faster for stimuli that were previously presented recently as opposed to more remotely in the past. This relationship between performance and presentation lag has been taken to reflect that memories are accessed by serially searching backwards in time, such that RT indicates the self-terminating moment of such a process. Here, we investigated the conditions under which this serial search gives way to more efficient means of retrieving memories. Event-related potentials (ERPs) were recorded during a continuous recognition task in which subjects made binary old/new judgments to stimuli that were each presented up to four times across a range of lags. Stimulus repetition and shorter presentation lag both gave rise to speeded RTs, consistent with previous findings, and we novelly extend these effects to a robust latency measure of the left parietal ERP effect associated with retrieval success. Importantly, the relationship between repetition and recency was further elucidated, such that repetition attenuated lag-related differences that were initially present in both the behavioral and neural latency data. These findings are consistent with the idea that a serial search through recent memory can quickly be abandoned in favor of relying on more efficient ‘time-independent’ cognitive processes or neural signals.

## 1. INTRODUCTION

Our mental representation of time and how we use it are important aspects of a variety of forms of cognition. Cortical circuits widely distributed throughout the brain computationally incorporate time into a wide array of decisions and behaviors, scaling across multiple orders of magnitude (e.g., Brown, Neath, & Chater, 2007; Howard, Shankar, Aue, & Criss, 2015; Ivry & Spencer, 2004; Paton & Buonomano, 2018). Aside from the low-level timing often involved in sensory and motor processing, time is also functionally coupled with our memory system. In the context of long-term memory, the influence of time has been shown in various situations, from distinguishing remote versus recent memories to providing a subjective (‘autonoetic’) awareness that allows for mentally traveling back and forth in time (Tulving, 2002; also see Murdock, 1974; Friedman, 1993). In the current study, we employed a continuous recognition procedure to investigate the behavioral and electrophysiological correlates of searching through time during memory retrieval. We demonstrate that, even over the course of seconds to minutes, there is a tradeoff between initially relying on time-related retrieval processing and the subsequent use of more efficient, thresholded (‘time-independent’) processes.

An experimental paradigm often used to assess the role of time in memory retrieval is the judgment of recency (JOR) task. In a basic version of this task, subjects are shown a list of successive stimuli to encode into memory and, following a delay, are presented with a pair of test stimuli from the list (e.g., Yntema & Trask, 1963; Morton, 1968). The task is then to choose the stimulus that occurred more recently (later in the list). When assessing such relative JORs, the standard finding is that judgments for more recently presented stimuli elicit shorter response times (RTs), suggesting that the past is searched sequentially and backwards, starting with the most recent stimulus (Muter, 1979; Hacker, 1980; also see Brown et al., 2007; Howard, 2018; cf. Chan, Ross, Earle, & Caplan, 2009). Critically, RTs have also been shown to vary only as a function of the more recent probe of the test pair, while being largely invariant to the time of the less recent probe. This result has been interpreted to reflect a self-terminating search process (Muter, 1979; Hacker, 1980; Hockley, 1984; McElree & Dosher, 1993), such that once the correct probe has been identified, the alternative becomes irrelevant to the judgment.

A simple interpretation of the findings described above is that there is a one-to-one correspondence between time and the behavioral outcome of the retrieval search process. However, at least two aspects of these and related empirical findings indicate that objective time differs from how it is represented mentally or the way in which that representation is employed in the context of memory retrieval. First, behavioral studies employing the JOR task have additionally shown that the relationship between behavioral measures and recency is nonlinear (e.g., Hinrichs, 1970; Hacker, 1980; Hockley, 1982). In an extension of Hacker’s (1980) classic study on short-term recency effects, Singh and Howard (2017) recently demonstrated that RTs were better fit, relative to a linear function, by a logarithmic (base 2) function of the recency of the correct probe. These replicated findings were taken as support for the idea that the memory search operates along a compressed timeline (Bjork & Whitten, 1974; Crowder, 1976) and are consistent with neural and modeling evidence that the internal representation of time exhibits properties of invariance across different timescales (for reviews, see Brown et al., 2007; Howard et al., 2015; Howard, 2018).

The second aspect of findings suggesting that time is not represented in a straightforward manner is related to an implicit assumption that is often made in studies of long-term memory. In particular, outside the domain of JORs which focus retrieval directly on time, and beyond short-term memory paradigms in which serial position (i.e. primacy and recency) effects dominate, the majority of retrieval task procedures disregard time as a relevant factor. For instance, in tests of recognition memory, encoding episodes are usually treated as though there is uniformity across time (within the encoding list), and behavioral measures associated with retrieval are consequently assumed to reflect a homogenous distribution. A particularly consistent treatment of this uniformity comes from studies that use electrophysiological measures such as event-related brain potentials (ERPs) to investigate memory retrieval. The resulting neural correlates are often evident within about a half second following stimulus onset, as is the case for the left parietal positivity for old relative to new test stimuli (for reviews, see Friedman & Johnson, 2000; Mecklinger, 2000; Rugg & Curran, 2007). The fact that these correlates of retrieval success occur within a shorter time frame than typical recognition RTs suggests that there is a disconnect between the neural signature of retrieval success and the behavioral correlate of terminating the backwards search. It remains to be determined whether neural measures such as ERPs, which are sensitive to the early processes contributing to memory retrieval, are sensitive to modulation according to the serial search across time.

To accommodate the factors mentioned above, continuous recognition tasks are particularly ideal. In these, subjects are presented with a single series of stimuli as opposed to the stimuli being segregated into distinct encoding and retrieval phases (Shepard & Teghtsoonian, 1961). Within the series, some stimuli are presented for the first time (new) and others are repeated (old), with a basic variant of the behavioral task being to make ongoing recognition (old vs. new) judgments. One advantage of these tasks, which is relevant to our interest here in the effects of time on memory, is that the lag between presentations can be manipulated from successive presentation (i.e. lag = 0) to any desired maximum. Several classic studies have shown that behavioral performance diminishes—in the form of both lower accuracy and longer RTs—as the lag between presentations increases (e.g., Shepard & Teghtsoonian, 1961; Donaldson & Murdock, 1968; Hintzman, 1969; Okada, 1971; Hockley, 1982; Friedman, 1990a), consistent with the JOR evidence and with the interpretation that access to less recent memories takes longer and becomes more likely to fail. Additionally, Singh, Oliva, and Howard (2017) recently demonstrated that recognition-related RTs increased logarithmically with lag, supporting a compressed representation of time over a matter of seconds to minutes.

In addition to the consistent effects of recency on retrieval, there is an extensive body of work on repetition affecting the retrieval process. Early studies showed that behavioral performance in the form of accuracy and RT was enhanced with further repetitions of study episodes (e.g., Hintzman, 1969; Hockley, 1982). These improvements have carried forth to the results of studies employing continuous recognition (e.g., Graetz, Daume, Friese, & Gruber, 2018; Johnson, Muftuler, & Rugg, 2008; Singh et al., 2017; Van Strien, Hagenbeek, Stam, Rombouts, & Barkhof, 2005). A common explanation of these findings is that repetition increases the strength of the memory trace (Morton, 1968; Murdock, Smith, & Bai, 2001), thereby making it more accessible to the retrieval search. This account thus provides an avenue for investigating the prevalence of the backwards serial search across time, which would presumably be replaced by more efficient, threshold-based signals operating on traces of different strengths. Contrary to this account, however, the aforementioned study by Singh et al. (2017) reported that the relationship between lag and RT did not diminish with an additional repetition of continuous recognition stimuli (also see Hockley, 1982). Thus, even though old stimuli were presented again for a second time, subjects still appeared to access their memories via the thorough, serial search process.

The purpose of the current study was to further examine the nature of the memory search process across time in continuous recognition, particularly in the light of additional stimulus repetitions. To this end, we manipulated the lag between presentations of a given stimulus to investigate the notion that the retrieval search operates along a continuous and compressed representation of time. Our design also involved presenting stimuli up to four times, in contrast with the three presentations used by Singh et al. (2017), enabling us to test whether further repetitions may be necessary to increase memory strength (i.e. fluency) to the point of overriding the serial search. Thus, one of our main predictions was that the relationship between RT and recency would diminish with additional repetition. Moreover, our experimental design was used in combination with ERPs, from which we assessed the latency of the left parietal positivity to characterize how the neural signature of retrieval success scaled with the presumed termination of the backwards serial search. Whereas prior ERP studies found no effect of lag on the peak latency of this ERP correlate (Friedman 1990a, 1990b; Rugg & Nagy, 1989), we make use of more robust methods of percent-area latency (for discussion, see Luck, 2014; Liesefeld, 2018) to provide evidence to the contrary.

## 2. METHODS

### 2.1. Participants

Nineteen subjects were recruited from the University of Missouri (MU) undergraduate subject pool and received partial course credit for participation. Informed consent was obtained in accordance with the MU Institutional Review Board. All subjects were right-handed, native-English speakers who had normal or corrected vision and no history of neurological disorders. The data from four subjects were removed from all analyses, due either to technical errors in stimulus presentation (N = 2) or having excessive artifact in the EEG (N = 2). The final sample of 15 subjects (7 females and 8 males) were between 18 and 24 years of age (*M* = 19). Of these 15 subjects, two subjects were missing data from one block of the task, and one subject was missing data from two blocks, all due to a technical error in data collection. Each of these subjects, however, had sufficient trial numbers according to the criteria described below.

### 2.2. Stimuli and Procedure

The stimulus pool consisted of 398 common objects in both color picture and word forms. Pictures subtended vertical and horizontal angles of about 3.1° each. Words were shown in white uppercase Arial font (approx. .7° × 4.1° for the longest word). All stimuli were presented on a solid gray frame (4.6° × 4.6°) at the center of the black background of a 24-inch widescreen LCD monitor (cropped to 1024 × 768 resolution), viewed at a distance of approximately 1 meter. The *Cogent 2000* toolbox (v. 1.32; http://www.vislab.ucl.ac.uk) was used to control stimulus presentation in MATLAB (v. R2012a; The MathWorks, Natick, MA).

Each subject completed six blocks of a continuous recognition task, with three blocks comprising only pictures and the other three comprising words. Picture and word blocks were presented in an ABBAAB order, with the starting type alternating across subjects. Each block used a random selection of 60 stimuli from the pool, with the remaining 38 stimuli used in a practice phase. Stimuli within a block were presented between one and four times, hereafter referred to respectively as *New, Old1, Old2*, and *Old3*. The repeated presentations resulted in 105 trials per block, and trials were organized into a series of 7 sub-blocks that were viewed by subjects as a continuous series of stimuli. The use of sub-blocks provided several features to the design: (1) it established a lead-in period, consisting primarily of new trials, to establish the initial presentation of stimuli that could eventually be presented multiple times; (2) it resulted in roughly a 1:1 ratio of new and old trials for most of the block (via the inclusion of new filler stimuli that were not analyzed); and (3) it allowed for systematic control over the lag between presentations of a given stimulus. Lag was defined as the number of intervening trials between a stimulus and its previous presentation (e.g., for a lag of 4, there were 3 intervening trials) and ranged from 4 to 35. On each trial, the picture or word stimulus was presented for 500 ms and followed by a central, white fixation marker (plus sign) for 1500 ms. Subjects indicated on each trial whether the stimulus being shown was a repetition (regardless of Old1, Old2, or Old3) or was novel (New) by respectively pressing the ‘.’ and ‘/’ keys on the keyboard with their right index and middle fingers.

### 2.3. EEG Acquisition and Processing

EEG was continuously recorded during all recognition blocks using a BrainAmp Standard system (Brain Vision LLC, Durham, NC) and elastic caps embedded with 59 Ag/AgCl ring electrodes (Easycap, Herrsching, Germany). Electrode locations were based on the International 10-20 system and included the following anterior/posterior chains of sites (from left to right): Fp1/z/2; AF7/3/z/4/8; F7/5/4/1/z/2/4/6/8; FT7, FC5/3/1/2/4/6, and FT8; T7, C5/3/1/z/2/4/6, and T8; TP7, CP5/3/1/z/2/4/6, and TP8; P7/5/3/1/z/2/4/6/8; PO7/3/z/4/8; and O1/2. Data were recorded with reference to an electrode placed at FCz, and the ground electrode was embedded in the cap at FT10. Additional electrodes were adhered to the mastoids (M1/2) for offline re-referencing, and below the left eye (IO1) and on the outer canthi (LO1/2) to capture vertical and horizontal EOG. Before the start of the first block, electrodes were manually adjusted until impedances were below 5 kΩ. The data were recorded at a 1-kHz sampling rate using an amplifier bandwidth of .01-100 Hz.

Processing of the EEG data was carried out with the *EEGLAB* toolbox (v. 14.0.0; Delorme & Makeig, 2004) in MATLAB. The continuous data were re-referenced to the mastoid average, down-sampled (200 Hz), band-pass filtered (.05-50 Hz), epoched (−500 to 1500 ms relative to stimulus onset), and baseline corrected with the pre-stimulus amplitude. Independent component analysis (ICA) was used to identify components indicative of eye-related artifacts (e.g., eye movements and blinks), which were rejected on the basis of high correlation with the electrode time courses. Epochs containing artifact were then manually rejected, a second pass of ICA was completed, and components seemingly related to additional artifacts (e.g., eye/muscle activity, noisy electrodes) were manually removed based on scalp topography and power spectra (Jung, Makeig, Westerfield, Townsend, Courchesne, & Sejnowski, 2000). The across-subject mean numbers of epochs included for the presentation conditions are as follows: New = 111.4 (range: 83-125), Old1 = 86.1 (65-101), Old2 = 78.9 (57-88), and Old3 = 65.4 (46-71).

### 2.4. Analyses

All of the analysis scripts for the results reported here, along with the relevant behavioral and EEG data, are available at https://osf.io/572jt/. The behavioral and EEG analyses were conducted in R (v. 3.5.0), using the *brms* (Bürkner, 2017), *BayesFactor* (Morey & Rouder, 2018), *bridgesampling* (Gronau & Singmann, 2017), and *lme4* (Bates, Martin, Bolker, & Walker, 2015) packages. For the EEG data, in addition to using EEGLAB for processing, the *latency* toolbox (Liesefeld, 2018; in MATLAB) was used for analysis, and the *MNE* package (Gramfort et al., 2014; in Python v. 3.6.4) was used for plotting purposes.

Prior to analysis, the response time (RT) data were checked for outliers, defined as >3 standard deviations from the z-scaled mean. Overall, 2.1% of trials (378 identified out of 18,344 data points) were excluded with this criterion. Trial-level measures (e.g., accuracy, RT, and ERP amplitude) were analyzed using a Bayesian multilevel modeling approach. A common model building procedure was used in which the maximal model (including all effects) was fit, effects were then randomly pruned, and the simplest model was retained in accordance with stabilized improvements of fit (also see Matuschek, Kliegl, Vasishth, Baayen, & Bates, 2017). All models used random intercepts for individual subjects and weakly informative priors [β ∼ *N*(0,1); σ ∼ *Cauchy*(0,2)]. Models were based on 2000 samples (the first half being warm-up samples) for each of four separate chains, providing adequate effective sample sizes and good convergence on the split-chain scale reduction factor (all *Rhat* ≤ 1.01).

Model comparison proceeded by comparing omnibus effects to an intercept only model and calculating Bayes Factors (*BF*s) via the bridge sampling technique. Following prior guidelines (Rouder, Engelhardt, McCabe, & Morey, 2016), ratios of *BF*s from different models were used to assess overall evidence for the inclusion/exclusion of interactions and main effects. *BFs* for non-nested models were calculated following Morey and Rouder (2018). Results are provided in terms of *BF*_*10*_, where values >1 refer to evidence favoring the alternative hypothesis (e.g., *BF*_*10*_ = 3 indicates the data are 3 times more likely to occur under the alternative than null) and positive values <1 refer to evidence in favor of the null (e.g., *BF*_*10*_ = 1/3 indicates data 3 times more likely to occur under the null). While our statistical interpretations are primarily based on these Bayesian estimates, frequentist statistics are also presented for comparison and to provide multiple estimates for evidence evaluation (Valentine, Buchanan, Scofield, & Beauchamp, 2019).

## 3. RESULTS

### 3.1. Behavioral Results

The mean proportions of correct responses on the continuous recognition task are shown in Figure 1A. As in our previous study with a similar behavioral design (Johnson et al., 2008), there was an apparent drop in proportion correct going from the first to second stimulus presentation (New to Old1), but performance then returned to a near-perfect level. An initial multilevel analysis of these data included predictor variables corresponding to repetition condition (New, Old1, Old2, and Old3) and stimulus material (pictures and words). Inclusion of the material predictor did not improve model fit, such that a model including only the repetition factor was preferred [*F*(1, 5527.8) = 0.03, *p* = .87, *BF*_*10*_ = 5.73e^-3^]. Next, an analysis including predictors for repetition and the lag between a stimulus and its prior presentation (range: 4-35, *M* = 16.8, *SD* = 6.8) was conducted for the proportion data. This analysis revealed strong evidence for a main effect of repetition [*F*(2, 3759.2) = 82.63, *p* < .001, *BF*_*10*_ = 9.84e^+30^] but evidence against both an effect of lag [*F*(1, 3760) = 2.45, *p* = .12, *BF*_*10*_ = 5.75e^-209^] as well as the interaction between lag and repetition [*F*(2, 3759.9) = 0.24, *p* = .79, *BF*_*10*_ = 2.84e^-4^]. Probing the repetition effect further with pairwise *t*-tests indicated lower accuracy for the Old1 compared to New, Old2, and Old3 conditions (all *p*s < .001; *BF*_*10*_ = 1.56e^+21^, 1.46e^+16^, 1.55e^+20^, respectively). Additionally, there was also evidence of an invariance between New and Old2 accuracy (*p* = .92, *BF*_*10*_ = 0.042) but insufficient evidence for or against differences involving the remaining comparisons (New vs. Old3: *p* = .01, *BF*_*10*_ = 1.25; Old2 vs. Old3: *p* = .01, *BF*_*10*_ = 0.893).

**Figure 1.**
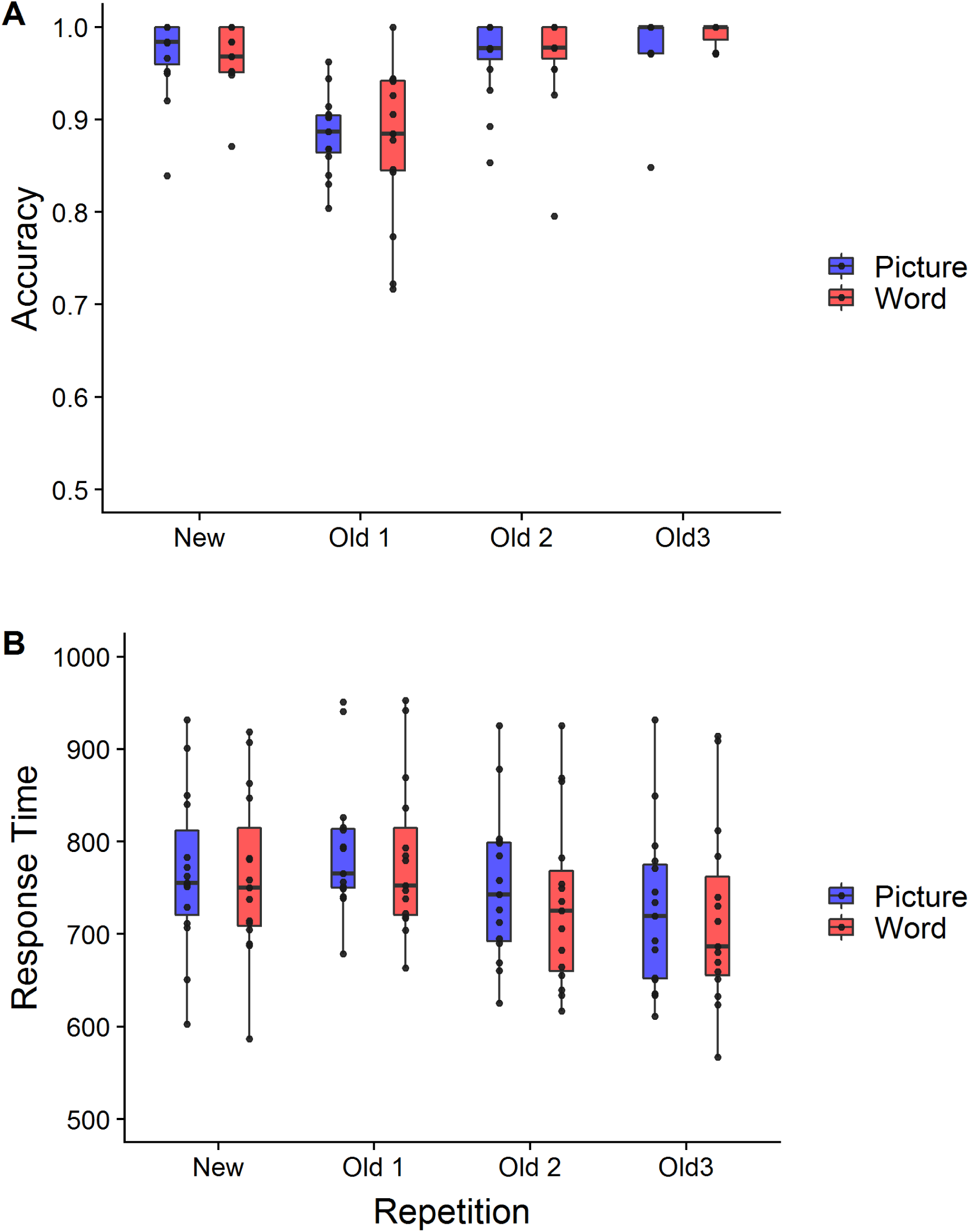
(A) Proportions of accurate recognition responses and (B) associated response times (RTs; in ms) for each repetition condition (New, Old1, Old2, and Old3) and stimulus material (pictures and words). Boxes indicate the median, 25th percentile, and 75th percentile; whiskers extend to the furthest value from the interquartile range (IQR) with a 1.5*IQR restriction; and individual dots represent subject measures.

The response times (RTs) corresponding to correct recognition judgments are summarized in Figure 1B. These data were first assessed according to whether changes due to lag occurred in a logarithmic manner, as has been demonstrated previously (Singh et al., 2017; also see Hinrichs, 1970; Hacker, 1980; Hockley, 1982). A modeling procedure that directly compared predictors corresponding to raw lag versus the logarithm (base 2) of lag indicated a better fit of the latter (ΔBIC = 5, *BF*_*10*_ = 1.85e^+4^). Due to this improvement, the log-transform of lag, hereafter referred to as “lag” for simplicity, was used in all remaining analyses. As with the analysis of correct proportions, there was no clear evidence for or against a material (picture vs. word) effect over just the effect of repetition, *F*(1, 3522.4) = 3.11, *p* = .08, *BF*_*10*_ = 1.07. Examining trial-level RTs in relation to the repetition and lag predictors provided support for a main effect of repetition [*F*(2, 3522) = 75.68, *p* < .001, *BF*_*10*_ = 4.62] and, to a lesser extent, a lag effect [*F*(1, 3522.1) = 29.93, *p* < .001, *BF*_*10*_ = 1.63]. Critically, though, there was strong evidence that lag interacted with repetition [*F*(2, 3520.2) = 4.81, *p* = .01, *BF*_*10*_ = 6.98e^+6^], an effect we further explore below.

To further understand the lag × repetition interaction, Figure 2 depicts the subject-and group-wise lag effects across repetition conditions. As is apparent, the lag effect is positive for the Old1 condition and attenuated for each additional repetition. The regression output from a model including this interaction of interest indicated that RTs in the Old1 condition increased with increasing lag at a rate of about 32 ms per doubling of lag [*b* = 31.80, 95% Highest Density Interval (HDI_95%_) [20.04, 42.87], *t*(3520.14) = 5.51, *p* < .001]. Moreover, this effect was weaker for the additional repetitions (Old2: *b* = 18.73, HDI_95%_ [-10.40, 46.68]; Old3: *b* = 3.34, HDI_95%_ [−26.95, 32.97]). Importantly, there was strong evidence for attenuation of the slope going from Old1 to Old3, *b* = −28.46, HDI_95%_ [−46.99, −9.90], *t*(3520.10) = −3.10, *p* = .002. There was also evidence, albeit weaker, for attenuation of the slopes going from Old1 to Old2, *b* = −13.07, HDI_95%_ [−30.44, 3.81], *t*(3520.26) = −1.50, *p* = .13, and from Old2 to Old3, *b* = −15.52, HDI_95%_ [--33.83, 3.26], *t*(3520.39) = 1.62, *p* = .11. Overall, these findings support our main prediction that multiple repetitions of old stimuli diminish the effects of recency (i.e. lag) on behavior.

**Figure 2.**
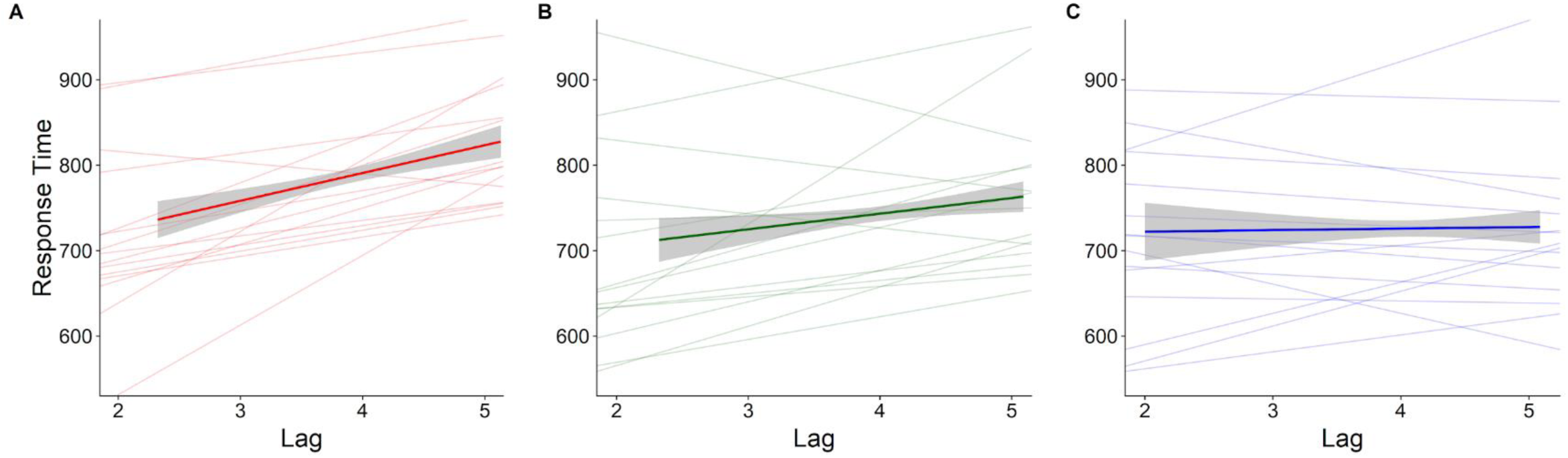
The relationship of log-transformed (base 2) lag between successive stimulus presentations and response time (RT; in ms), plotted separately for the (A) Old 1, (B) Old 2, and (C) Old 3 conditions. Regression lines for individual subjects are thin, whereas the mean regression lines (and 95% confidence bands around the mean) are thick.

### 3.2. ERP Results

Analysis of the ERPs first focused on characterizing the left parietal old/new effect (for reviews, see Friedman & Johnson, 2000; Rugg & Curran, 2007) for each of the repetition conditions. Next, we sought to test for differences in the latency of this effect with respect to repetition, analogous to the main effect of repetition observed for the RT data. Finally, to provide converging neural evidence for the lag-related differences identified behaviorally, we split the trials into two groups according to lag and tested for differences in the latencies of the corresponding old/new ERP effects.

Figure 3A displays the grand-average ERPs for a representative set of nine electrodes covering the scalp. As shown, the ERPs for the Old1, Old2, and Old3 conditions appeared more positive-going than those for the New condition. These effects onset by about 300 ms after stimulus onset and lasted for another 500-700 ms. Additionally, as is apparent from the topographic scalp maps shown in Figure 3B, the old/new differences were widespread, ranging over bilateral scalp and from frontal to parieto-occipital electrodes.

**Figure 3.**
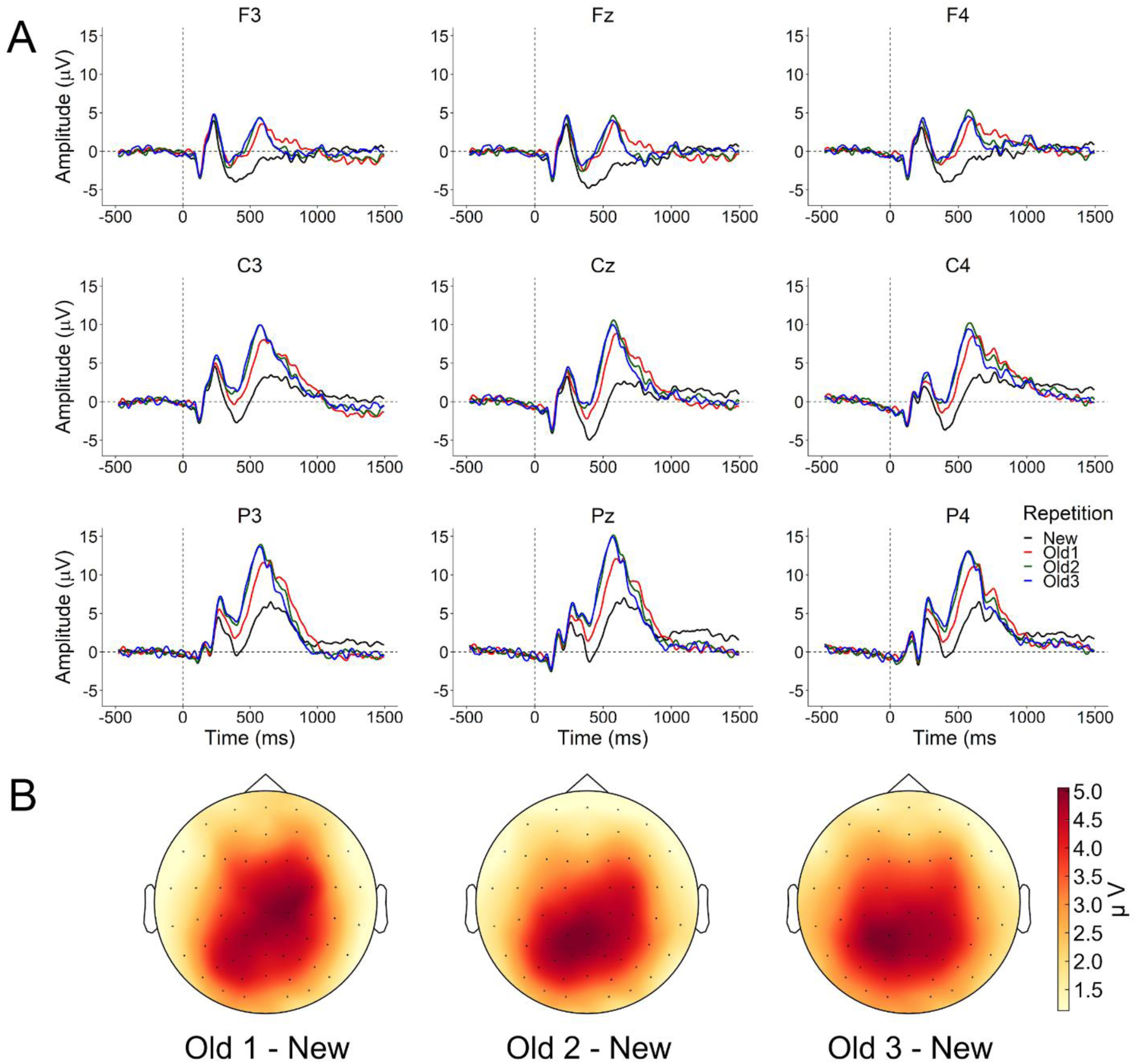
(A) Grand-average ERPs for the repetition conditions (New, Old1, Old2, and Old3) from a representative set of nine electrodes (F3/Fz/F4, C3/Cz/C4, P3/Pz/P4) across the scalp. (B) Topographic scalp maps depicting differences between the Old1/Old2/Old3 and New conditions.

To further characterize the old/new ERP effects, and to examine any modulation of these differences according to repetition, the average amplitudes between 500-800 ms post-stimulus onset were extracted from the montage of nine electrodes shown in Figure 3A. Multilevel modelling using factors of repetition (New, Old1, Old2, and Old3), anterior/posterior chain (frontal, central, and parietal), and laterality (left, midline, and right) gave rise to a main effect of repetition [*F*(3, 46125) = 566.01, *p* < .001, *BF*_*10*_ > 1.0e^999^] and an interaction between repetition and anterior/posterior chain [*F*(6, 46093) = 9.13, *p* < .001, *BF*_*10*_ = 1.0e^18^]. This interaction can be described, as is apparent in Figure 3, in terms of the old/new differences being larger over central and parietal sites compared to frontal sites. Additionally, we tested a model that was restricted to data only from the old conditions (Old1, Old2, and Old3); strong evidence for the repetition effect remained, *F*(2, 31087) = 7.92, *p* < .001, *BF*_*10*_ = 14.38, indicating that the magnitude of the old/new differences changed with repetition (as opposed to the old conditions merely being distinguishable from the New condition). To further simplify these results and to increase the signal-to-noise ratio for the latency analyses reported below, the old/new ERP differences were averaged over a group six electrodes (CP3/CP1/CPz and P3/P1/Pz) where the effects were maximal. The resulting waveforms according to repetition condition are shown in Figure 4A.

**Figure 4.**
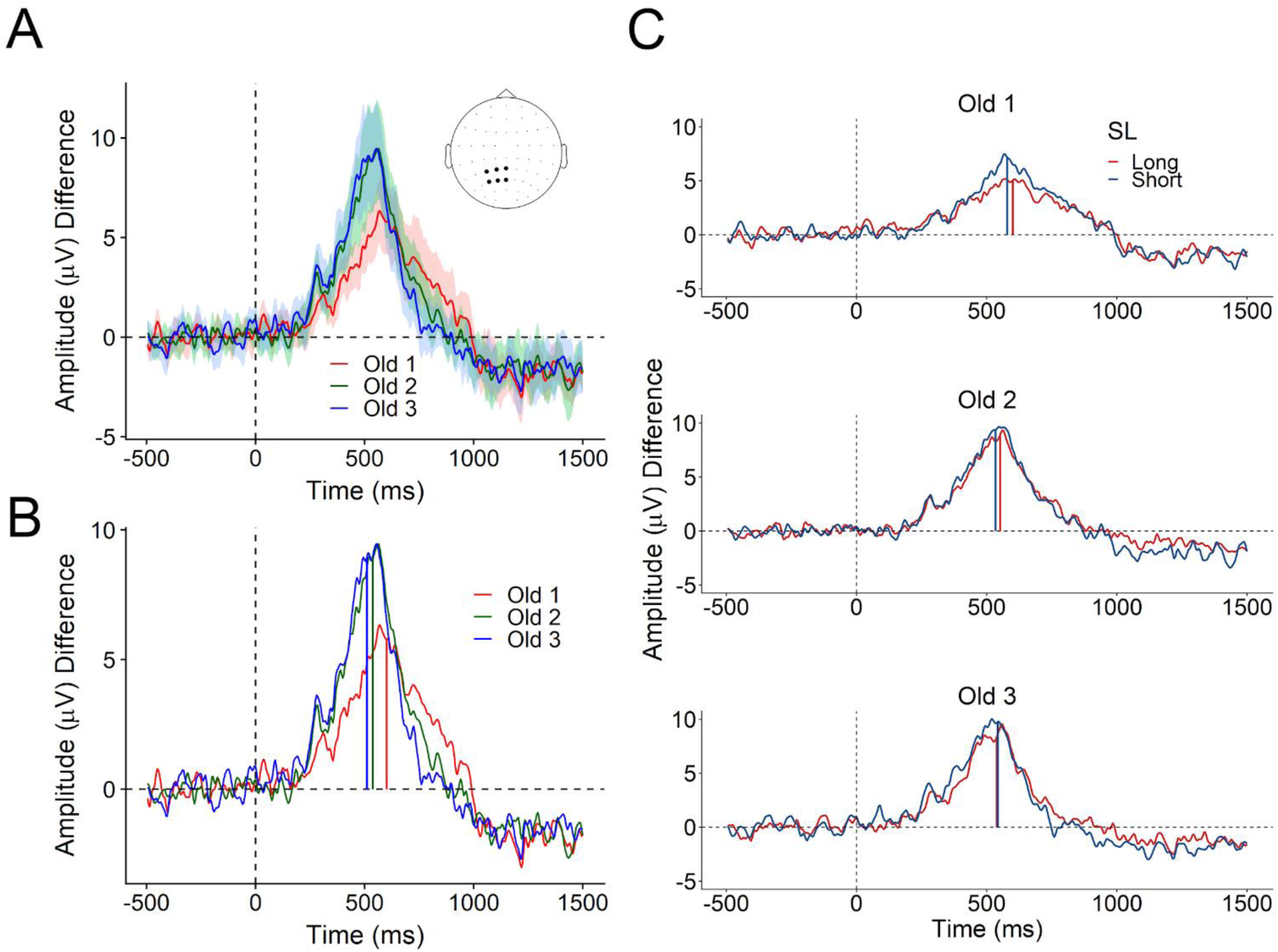
(A) Old/new ERP differences according to repetition condition (Old1, Old2, and Old3), averaged over a montage of the six electrodes from left posterior scalp (CP3, CP1, CPz, P3, P1, and Pz). The shaded bands indicate 95% confidence intervals around the mean. (B) Old/new ERP differences from left posterior scalp (as in Panel A), along with the percent-area latency estimate for each repetition condition shown by a vertical line. (C) Old/new ERP differences from left posterior scalp according to each repetition condition and the median split of lag (short and long). Vertical lines again indicate the percent-area latency estimates for short and long lags.

Having established differences in the amplitude of the left parietal old/new effect across repetition conditions, we next turned attention to identifying any differences in the latency of this effect. The timing of the effects was defined as the percent-area latency, which is the time at which the effect of interest (here, the old/new difference) reaches a specified percentage of its area under the curve. Percent-area latency has been shown to be more robust to various sources of noise than alternative measures such as peak latency or onset latency (Liesefeld, Liesefeld, & Zimmer, 2016; Liesefeld, 2018). We defined the latency measure by first establishing the onset and offset as 30% of the peak amplitude difference within our time window of interest (500-800 ms) and then determining the time at which the area under the curve reached 50% (also see Liesefeld, 2018). As shown in Figure 4B for the group of six left posterior electrodes (see above), when comparing the latencies of the old/new effects according to repetition, the effect for the Old1 condition appeared later than that for the Old2 and Old3 conditions. A one-way ANOVA of these measures confirmed a main effect of repetition on latency, *F*(2,28) = 17.18, *p* < .001, *BF*_*10*_ = 13.43. Pairwise comparisons revealed that latency differences were largest between the Old1 (*M* = 601 ms, *SD* = 95) and Old3 (*M* = 511 ms, *SD* = 50) conditions, *t*(28) = 5.70, *p* < .001, *BF*_*10*_ = 12.73. There was also positive evidence for a difference between the Old1 and Old2 (*M* = 537 ms, *SD* = 55) conditions (*t*(28) = 4.04, *p* = .001, *BF*_*10*_ = 2.16) but little evidence for a difference between Old2 and Old3 (*t*(28) = 1.66, *p* = .24, *BF*_*10*_ = 0.694).

Finally, to test specifically for differences in the latencies of the old/new ERP effects according to lag, the data for each repetition condition were split into two groups—hereafter referred to as *short* and *long* lag—relative to the median lag determined separately for each subject (*M* = 16.2, *SD* = .7, across subjects). The decision to split the data into two groups was based on maximizing the amount of data included in each group, as the trial-level ERP latencies were expected and confirmed to be too noisy for a regression-based approach. We note here that our design allowed for capitalizing on large numbers of trials, even after the split (*M* = 35.8 trials per repetition × short/long combination; range: 20-50 trials per cell). Figure 4C displays the resulting median-split ERPs for each repetition condition. As only the Old1 condition exhibited reliable differences in RT according to lag, we expected that only this condition would show lag-related differences in ERP latency. This was confirmed by a direct comparison of the lag conditions, revealing that the latency was earlier for short (*M* = 578 ms, *SD* = 75) relative to long (*M* = 600 ms, *SD* = 92) lags, *t*(14) = 2.09, *p* = .03, *BF*_*10*_ = 2.73. (Given the directional predictions afforded by the RT differences reported earlier, the alternative hypothesis for each of these tests was restricted to the negative portion of the prior distribution [*r* = .707, *-Inf < d < 0*].) By contrast, there was no evidence for any differences between short and long lags for either the Old2 (*M*s = 538 and 547 ms, *SD*s = 54 and 59, respectively; *p* = .27, *BF*_*10*_ = 0.455) or Old3 (*M*s = 533 and 536 ms, *SD*s = 63 and 55, respectively; *p* = .44, *BF*_*10*_ = 0.298). These findings of lag-related ERP latency differences for the Old1 condition, but not for the Old2 and Old3 conditions, thus provide converging evidence with the positive relationship observed between lag and Old1 RTs.

## 4. DISCUSSION

There has been a longstanding debate in human memory research between the idea that memory traces are organized along a mental representation of time (e.g., Muter, 1979; Hacker, 1980; Hockley, 1984; Howard, 2018) versus the notion that changes in memory performance reflect factors merely correlated with time, such as strength or fluency (e.g., Morton, 1968; Murdock et al., 2001). In experimental paradigms that focus memory retrieval on time, as is the case with those requiring judgments of recency (JORs) or manipulating the lag between stimulus repetitions during continuous recognition, RTs have regularly been shown to increase with the time passed since a stimulus’s prior occurrence (e.g., Hacker, 1980; Singh & Howard, 2017). This pattern of findings suggests that the retrieval search operates in a backwards, self-terminating manner (Howard, 2018). In the current study, a continuous recognition task was used to track such lag-related changes in behavior and in a robust measure of the latency of ERP correlates of retrieval success. Consistent with memory search operating along a timeline, we observed that the RTs on old trials and the corresponding latency of the left parietal old/new ERP effect were longer with increasing lag. Specifically, the lag effects were strongest for the first repetition of stimuli (our Old1 condition). Moreover, because the RT measures were stable enough to allow for trial-level Bayesian multilevel modeling, the present findings also provide support for the idea that the retrieval search in our continuous recognition task operates along a compressed (sublinear) representation of time (also see Singh et al., 2017; Howard, 2018).

In addition to providing a combination of behavioral and neural evidence supporting a time-based retrieval search, another goal of the current study was to determine how the likelihood of employing such a thorough, serial process might change with additional stimulus repetitions. The motivation for investigating this comes from a long history of studies indicating that retrieval can be based on highly-efficient processes (or signals) related to memory strength or fluency (Hintzman, 2005; Morton, 1968; Murdock et al., 2001). In particular, several continuous recognition studies have shown that repetition leads to enhanced performance in both accuracy and RTs (e.g., Graetz et al., 2018; Hintzman, 1969; Johnson et al., 2008; Singh et al., 2017; Van Strien et al., 2005). We replicate those findings here, as shown in Figure 1. However, as described in the Introduction, a recent study by Singh et al. (2017) showed that repeating stimuli an additional time (corresponding to our Old2 condition) did not affect the magnitude of the positive relationship between RT and lag, suggesting that the memory strength of those stimuli was not sufficient for overcoming the time-based search. In the current study, we further tested for such an effect by including a fourth presentation (Old3), which turned out to exhibit the largest attenuation, relative to the Old1 condition, in the slope of the lag effect on RT (cf. Singh et al., 2017). Furthermore, consistent with diminished slopes of RT effects, there were no differences (via Bayes Factor analysis) across the median split of lags for the latencies of left parietal old/new ERP effects for the Old2 and Old3 conditions. Together, these findings provide novel behavioral and neural evidence for the idea that the backwards search can be replaced by more-efficient processing related to strength or fluency. One possibility suggested by our findings is that a thresholded process takes over, as indicated by the similarity in both the amplitudes as well as the latencies of the ERP effects for the Old2 and Old3 conditions (see Figure 4).

While the task demands of the continuous recognition procedure used in the current study are essentially identical to those of Singh et al. (2017), we suspect that some other aspects of experimental design might have led to the different patterns of results. First, Singh et al. used a wider range of lags— up to 128—compared to the maximum of 35 in the current study. An obvious assumption can be made about the stimuli in the longest lag conditions, which is that they are likely associated with weak memory traces in comparison to those corresponding to shorter lags. Consequently, the inclusion of such long lags could have promoted a strategy of maintaining the thorough, backwards memory search, so as not to miss any of these relatively weak traces. By contrast, because our study used a range of shorter lags, the serial search may have become unnecessary for subjects to perceive that they were achieving a sufficient level of retrieval success.

Another design difference between the current study and the one by Singh et al. (2017) concerns the relative probabilities of old and new trials in the continuous recognition blocks. As discussed earlier, our study included an additional repetition (the Old3 condition) that was not present in the previous work, which thereby contributed to increasing the ratio of old to new trials. Even without these trials, there was still a substantial difference in the overall old/new ratio between the two studies, due to an inclusion of many more new stimuli throughout the Singh et al. study. In particular, old stimuli in the three experiments of that study respectively accounted for only 16%, 28%, and 20% of the total number of trials. In the current study, by comparison, we explicitly controlled (via the sub-block structure, as described in Stimuli and Procedure) the prevalence of old trials at about 50% for the majority of each block. Even when accounting for the initial lead-in period, which was dominated by new stimuli, the old conditions still accounted for 43% of trials. Just as the longer lags used by Singh et al. could have encouraged the serial search, our higher proportion of repeated stimuli could have led to a strategy of consistently relying on a more efficient, threshold-like property for making recognition judgments.

Although the theoretical frameworks we have thus far worked within the confines of here focus on time-versus strength-based retrieval processes, there are other alternative theories worth considering in regard to changes in memory traces that correlate with time. In particular, some of these theories posit that traces are not organized along a timeline *per se*, but can be thought of as part of a composite memory store (Anderson, 1973; Murdock, 1982; Shiffrin, Ratcliff, Nurnane, & Nobel, 1993). By such an account, stimuli occurring further in the past are subject to increased degradation (cf. fidelity) of trace quality, leading to diminished accessibility and thereby slower RTs. On the face, this interpretation is indistinguishable from those suggesting that recency modulates the strength (or fluency) of a memory (see Hintzman, 2005; Hintzman, 2016). The findings of the current study offer little in the way of enhancing any distinction, with the exception of one possibility. That is, the latency differences observed here exist for ERPs in addition to behavioral responses, thus pushing the lag-related effects on retrieval success to an earlier time windows after retrieval cue onset. The earlier that such latency effects can be demonstrated, the more likely they are to be indicative of the backward search process, as the composite account would instead predict that longer RTs and later-onsetting ERP correlates would reflect prolonged processing for degraded traces. One potential avenue to explore in future research, therefore, is to test whether ERP latency differences can be identified in the presence of equivalent RTs that have been taken as support of the composite account.

To conclude, in the current study we demonstrate further evidence, in the form of behavioral RTs, that both repetition and recency (lag) influence recognition judgments. We also extend those effects to the domain of ERPs in two ways. First, multiple stimulus repetitions increased the amplitude of the left parietal old/new effect (also see Graetz et al., 2018; Van Strien et al., 2005). Second, we provide evidence that the latency of the old/new ERP effect was shortened by repetition (for analogous results, see de Chastelaine, Friedman, Cycowicz, & Horton, 2009; Liesefeld et al., 2016; Park & Donaldson, 2016) as well as recency, further indicating the viability of using robust measurement approaches to understand the precise timing of cognitive processes (Liesefeld, 2018; Ouyang, Herzmann, Zhou, & Sommer, 2011; Smulders, 2010). Critically, the relationship between recency and the latencies of the behavioral and neural correlates was shown to be attenuated with additional presentations of recognized stimuli. While we cannot conclusively rule out alternative theoretical accounts that involve the storage of composite traces, our convergent behavioral and ERP findings are consistent with the idea that recent memories initially exhibit a form of temporal organization. However, as memories become more accessible, recognition judgments may shift away from a reliance on an ordered timeline to more efficient, and perhaps thresholded, cognitive processes or neural signals.

